# Epigenetic features of far northern Yakutian population

**DOI:** 10.1101/2023.07.19.549706

**Authors:** Alena Kalyakulina, Igor Yusipov, Elena Kondakova, Maria Giulia Bacalini, Cristina Giuliani, Tatiana Sivtseva, Sergey Semenov, Artem Ksenofontov, Maria Nikolaeva, Elza Khusnutdinova, Raisa Zakharova, Maria Vedunova, Claudio Franceschi, Mikhail Ivanchenko

## Abstract

Yakuts are one of the indigenous populations of the subarctic and arctic territories of Siberia characterized by a continental subarctic climate with severe winters, with the regular January average temperature in the regional capital city of Yakutsk dipping below −40°C. The epigenetic mechanisms of adaptation to such ecologies and environments and, in particular, epigenetic age acceleration in the local population have not been studied before. This work reports the first epigenetic study of the Yakutian population using whole blood DNA methylation data, supplemented with the comparison to the residents of Central Russia. Gene set enrichment analysis revealed, among others, geographic region-specific differentially methylated regions associated with adaptation to climatic conditions (water consumption, digestive system regulation), aging processes (actin filament activity, cell fate), and both of them (channel activity, regulation of steroid and corticosteroid hormone secretion). Further, it is demonstrated that the epigenetic age acceleration of the Yakutian representatives is significantly higher than that of Central Russia counterparts. For both geographic regions, we showed that epigenetically males age faster than females, whereas no significant sex differences were found between the regions.

## 1. Introduction

The Yakuts (Sakha) are people living in the subarctic and arctic territories of eastern Siberia. Anatomically modern humans inhabited the region of the modern Republic of Sakha about 30,000 years ago [1], moving from the west, where Sakha connects to the inner Eurasian steppe belt through southern Siberia [2, 3]. The northeastern part of Yakutia is located on one of the main migration routes from the southern regions of the Yenisei, Amur and Baikal coasts to the Arctic coast and to America [4]. Thus, the native inhabitants of Siberia, including Yakuts, are also viewed with regard to American colonization [5, 6]. There is evidence bringing Yakuts closer to Amerindians in terms of genetic variability, which may indicate a common ancestry of Siberians and Native Americans [7]. The most recent wave of migration to the territory of modern Yakutia involved the Tungus of Transbaikalia and Turkic-speaking Yakuts of the Western Baikal region, and took place about 2,000 years ago [8].

Eastern Siberia is one of the permanently inhabited regions with an extremely cold climate. From the evolutionary perspective, populations surviving in such an extreme climate for many years should have accumulated genetic changes adapting them to the cold and to other local factors such as seasonal extremes in daylight, food availability, etc. [9]. Indeed, low serum lipid levels were identified and related to the increased energy metabolism [10], and higher blood pressure was also observed [11, 12]. A large study of Siberian populations identified candidate genes for adaptation to the cold, associated with energy and metabolic regulation, as well as contraction of vascular smooth muscle [9]. Besides, leptin and irisin, which play an important role in the processes of adaptation to the cold, become the focus of several other studies of the Yakutian cohort [13–15]. However, genetic studies for Siberian populations are often limited to mitochondrial DNA, Y chromosome or single-nucleotide polymorphisms [8, 9, 16, 17], while epigenome-wide studies have not been performed.

Epigenome-wide association studies (EWAS) usually analyze DNA methylation and focus on differentially methylated CpG sites (differentially methylated positions or DMPs), similar to single-nucleotide polymorphisms in genome-wide association studies (GWAS) [18]. EWAS detects epigenetic differences between human cohorts by quantitatively comparing methylation levels of thousands of CpG sites [19]. Individual studies have shown that significant ethnic differences in DNA methylation are reflected in cell composition, risk of some noncommunicable diseases [20]; are associated with blood lipid levels [19], body mass index [21], liver function [22] and blood pressure [23]. There are also studies showing environmental [24, 25] and socio-economic [26–28] influence on DNA methylation. The influence of race/ethnicity has been also studied in the context of health status, mortality, and susceptibility to diseases [29, 30], as well as epigenetic aging speed [31, 32]. Different frequencies of certain genetic variants can lead to epigenetic differences between ethnicities [33–35] and contribute to specific patterns of epigenetic aging.

One of the best-known mechanisms for tracking changes in DNA methylation are epigenetic clocks that are models aggregating information about a limited set of CpG sites and used to construct estimates of epigenetic age and mortality risk. The most common clocks are Hannum DNAmAge [36], Horvath DNAmAge [37], GrimAge [38], DNAmPhenoAge [39]; the metric of accelerated epigenetic aging as a deviation of chronological age from that predicted by the clock is also often considered.

Another interesting aspect would be addressing sex specific differences in DNA methylation for the cohorts of different ethnicities that live in different climates. EWAS reported differences in DNA methylation associated with sex differences in genes on the autosomes [40, 41]. Moreover, regions of different methylation in males and females are not limited to the sex chromosomes but are genome-wide and can be tracked in various tissues, e.g., brain, pancreas, blood [42–44]. Epigenetic profiles undergo profound changes during aging, with differences between males and females either remaining [45, 46] or changing in different regions of the genome [40, 47]. It can be hypothesized that such sex-specific DNA methylation trajectories may contribute to sex differences in survival [48].

This work is the first epigenetic study focused on the Yakutian population. Since the distinctive feature of this ethnic group is living in severe climatic conditions, it can shed light on the epigenetic features of the adaptation mechanisms to the cold. Further, since severe climate can affect both health and aging, it is interesting to assess age acceleration and aging rate for the Yakutian population. Another aspect of the study is the sex-specific methylation patterns in the Yakutian cohort.

The whole blood DNA methylation data of the residents of Yakutia and Central Russia collected in this work allows us to perform a comparison between these two cohorts living in very different environmental and climatic conditions. The wide age range and the balanced number of representatives of both regions allows us to compare them and identify region-specific features of DNA methylation. Region-specific CpG sites can be detected by searching for differentially methylated positions (DMPs), and EWAS can analyze them in terms of the corresponding biological processes and pathways. Epigenetic clocks allow us to determine the age acceleration in the considered regions and compare the aging rate between them, as well as to estimate the blood cell composition. Similar age and quantitative distributions of males and females from Yakutia and Central regions in our data enable even deeper analysis and investigation of sex-specificity between the regions. Following the same pipelines as for the region-specificity analysis, DMPs between males and females in both regions can be found and analyzed using EWAS; epigenetic ages, age acceleration, blood cell composition can also be investigated for males and females. This could reveal whether there are sex differences specific to particular regions.

## 2. Results

### 2.1. Participants and study design

This study involved whole blood DNA methylation data from 245 healthy participants collected in 2020-2022 in the Central region of Russia (Nizhny Novgorod, Vladimir, and Moscow regions, highlighted yellow in Figure 1A) and Yakutia (Republic of Sakha, highlighted gray in Figure 1A). All participants from Yakutia are indigenous people, born and living in Yakutsk or in the nearby uluses (villages). The Central region includes 131 samples (78 females and 53 males) and the Yakutia region includes 114 samples (63 females and 51 males). Information about participants are presented in Supplementary Table S1. Figure 1B shows the age distributions in the two regions. In the Central region the age of participants ranged from 15 to 101 years, in the Yakutia region from 11 to 99 years.

**Figure 1.**
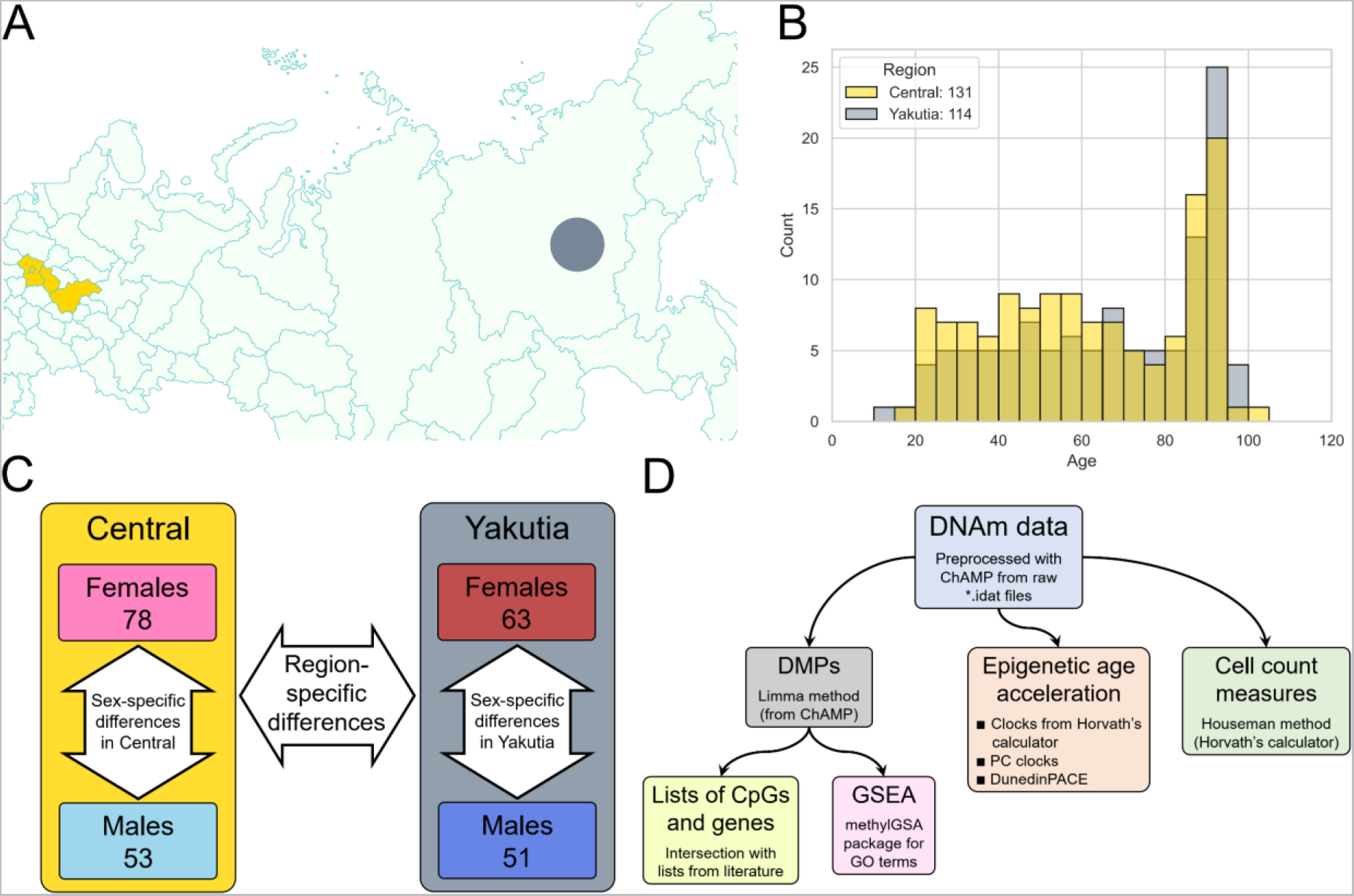
Participants and study design. (A) Part of the administrative map of Russia with the highlighted regions where participants were recruited for the study. Central region is highlighted in yellow; part of the Yakutia region is highlighted in gray. (B) Histogram of the age distribution in the two regions. (C) The scheme of EWAS experiments in this study: first, epigenetic differences between regions are studied, then sex-specific differences in both regions are sought and compared between the regions. (D) The basic workflow that the EWAS experiments follow. It involves searching for DMPs - CpGs that are statistically different between the phenotypes in question - which are used for Gene Set Enrichment Analysis (GSEA) and comparison with lists of CpGs and genes from other works. In addition, the analysis of various biomarkers obtained from DNAm data (epigenetic ages, their derivatives, and blood cell count measures).

The compared Central region of Russia and Yakutia differ significantly in their climatic conditions. The average winter temperature in Nizhny Novgorod is −13°C (most participants from the Central Russia live in Nizhny Novgorod and Nizhny Novgorod region), while Yakutia is one of the coldest geographical regions on Earth with a permanent population, the average winter temperature in Yakutsk reaches −42°C. The duration of the negative temperature period is also different: from November to March in Nizhny Novgorod and from October to April in Yakutsk. The difference in average temperatures between the warmest and coldest periods is extremely high in Yakutsk, it reaches 70°C, while in Nizhny Novgorod it is about twice less [49].

Life expectancy at birth in the Republic of Sakha is 69.98 years in 2021, somewhat exceeding the life expectancy in Nizhny Novgorod region, which is 68.93 years, despite the harsher climate. It is worth noting that in both regions the life expectancy of males and females differs significantly. Male life expectancy at birth in Yakutia in 2021 is 65.65 years, in Nizhny Novgorod region is 63.81 years; female life expectancy at birth in Yakutia in 2021 is 74.47 years, in Nizhny Novgorod region is 73.97 years [50]. The average age differs significantly: 35.0 years in Yakutia and 42.9 years in Nizhny Novgorod region. Significant differences are also observed between the sexes: the average age of males in Yakutia is 33.2 years, in Nizhny Novgorod region is 39.7 years; the average age of females in Yakutia is 36.6 years, in Nizhny Novgorod region is 45.5 years.

DNA methylation data for 245 samples were collected using Illumina Infinium MethylationEPIC BeadChip technology that measures DNA methylation levels from a total number of 866,836 genomic sites with single-nucleotide resolution. After all preprocessing procedures, 739,168 CpG sites remained.

EWAS was performed in three different settings (Figure 1C): EWAS between two regions, Central Russia and Yakutia (Section 2.2), as well as two EWAS investigating sex differences in these two regions (Section 2.3). The results of these two sex-specific studies were also then compared with each other.

Each EWAS study followed the workflow shown in Figure 1D. Differentially Methylated Positions (DMPs) analysis allows to perform a contrast comparison between two phenotypes for the considered covariate (in our case, region or sex, depending on the task). The top of the most significant CpG sites and corresponding genes were considered: enrichment of chromosomes, genomic regions, and CpG islands were studied, and overlaps with similar lists of specific CpG sites and genes from published works were examined. The resulting FDR-corrected [51] p-values of the statistical test for the difference in methylation levels between the two considered groups for all CpGs were used to perform Gene Set Enrichment Analysis (GSEA). The resulting significant terms from the Gene Ontology (GO) library [52, 53] were then analyzed and interpreted from a biological perspective.

Next, methylation data were uploaded to Horvath’s online calculator [54] - it allowed to obtain values for the 4 most common estimators of epigenetic age: Horvath DNAm age [37], Hannum DNAm age [36], DNAm PhenoAge [39], and GrimAge [38] and also blood cell count measures for the following cell types: CD8T, CD4T, NK, Bcell, Mono, Gran, according to Houseman algorithm [55]. In addition to the classical versions of epigenetic clocks, we considered their retrained with principal components variations, which significantly increase the reliability of classical epigenetic models [56]. Another epigenetic biomarker used in the work, which characterizes the rate of aging, is DunedinPACE - estimation of the rate of aging by DNA methylation. This metric can be interpreted as the number of biological years per chronological year (with a mean value of 1) [57].

### 2.2. Region-specific differences

In this section, epigenetic differences between the two regions - Central Russia (131 samples) and Yakutia (114 samples) - are analyzed according to the workflow shown in Figure 1C.

#### 2.2.1. Region-specific CpGs, DMPs, and GSEA

To investigate the epigenome-wide differences between the two regions, we first applied the limma method [58] to search for region-specific CpGs (DMPs). Figure 2 shows the results of this algorithm.

**Figure 2.**
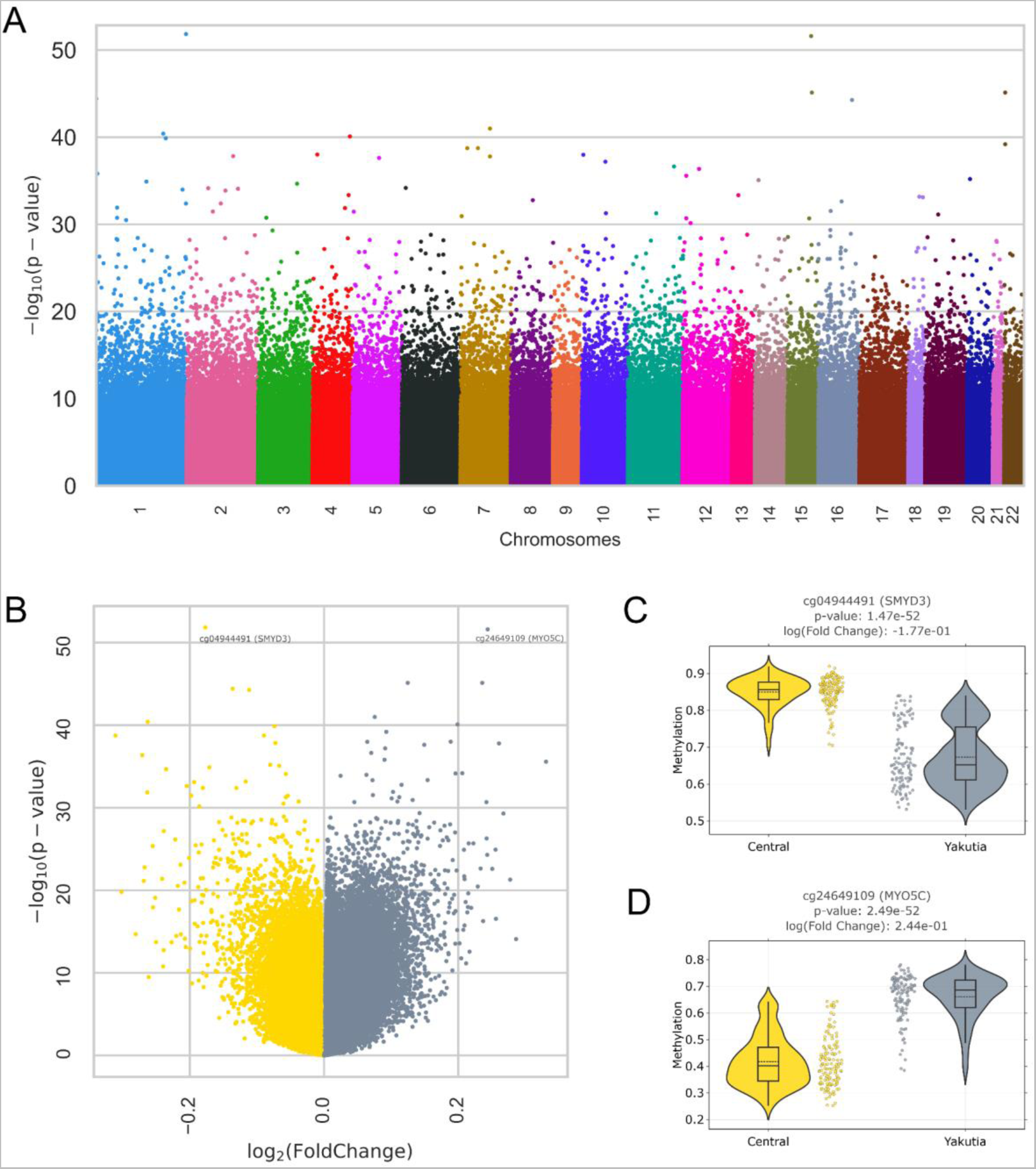
DMP analysis (limma) of regional specificity. (A) Manhattan plot for the distribution of corrected p-values of region-specific CpGs distributed by location in chromosomes; (B) Volcano plot of limma results for all CpGs. Hypermethylated CpGs in the Central region are highlighted in yellow, and hypermethylated CpGs in Yakutia are highlighted in gray. The two most significant CpGs are highlighted in the plot; the distributions of methylation levels between regions are shown in (C) cg04944491 (SMYD3 gene) and (D) cg24649109 (MYO5C gene). The solid line on the boxplot corresponds to the median value; the dotted line corresponds to the mean value.

The Manhattan plot (Figure 2A) illustrates the ordering by chromosomal position of CpG sites with different statistical significance (FDR-corrected p-value) in the methylation level between the two regions. Volcano plot (Figure 2B) in addition to statistical significance contains absolute value of logarithmic fold change, which indicates relative methylation level: yellow dots with negative log2(FoldChange) values correspond to CpG-sites hypermethylated in Central region relative to Yakutia (Figure 2C), whereas gray dots with positive log2(FoldChange) values are hypermethylated in Yakutia region relative to Central region (Figure 2D).

We identified the Top-1000 CpG sites with the most statistically different methylation levels between regions with the lowest p-values (Supplementary Table S2) and considered the distribution and enrichment analysis of these CpGs for chromosomes, genomic regions, and CpG islands (Figure 3).

**Figure 3.**
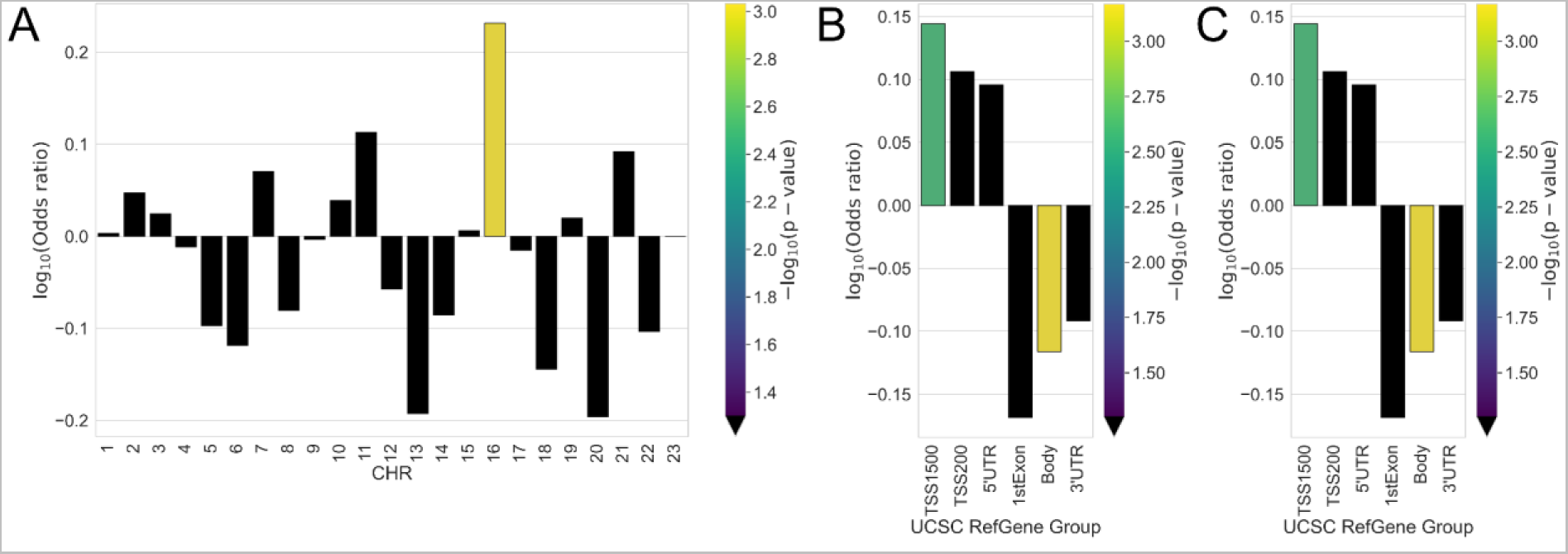
Enrichment of (A) chromosomes, (B) genomic region, and (C) relation to CpG island for Top-1000 region-specific CpGs. Odds ratio values and corresponding p-values (shown by color) were obtained from Fisher exact test. Black color indicates the absence of statistical significance (p-value > 0.05).

These region-specific probes were statistically significantly overrepresented only in chromosome 16, and no statistically significant features were detected in all other chromosomes. Overrepresentation of such CpGs was also found in the regulatory genomic region TSS1500 and underrepresentation in the gene body. It is noteworthy that the selected Top-1000 CpGs belong mainly to islands, regions with increased CpG density (overrepresentation with p-value < 1e-25), CpGs in OpenSea (regions with the lowest density) are statistically underrepresented. Since one of the main functions of DNA methylation is to regulate gene expression, a large number of CpG islands are located in the promoter regions of the gene [59], which is also reflected in the obtained result.

Interesting that chromosome 16 contains numerous erythroid-specific alpha-globin genes, and hypermethylation of certain differentially methylated regions at different stages of erythropoiesis has been shown [60]. In the context of differences between the considered regions, it would be interesting to consider the relationship between red blood cell count (RBC) and adaptation to cold temperatures; however, very few such studies have been performed. In [61] it was shown that exposure to extremely low temperatures for 50 days can lead to a decrease in RBC.

It is interesting to compare the obtained Top-1000 CpGs, corresponding to 656 genes, with the list of 791 genes from the work Cardona et. al. [9], analyzing single-nucleotide polymorphism data for 10 indigenous Siberian populations, thereby describing the genetic influence/impact on epigenetic variation. These 791 genes are associated with biological processes and pathways hypothesized to be involved in cold adaptation and related to basal metabolic rate, non-shivering thermogenesis, response to temperature, smooth muscle contraction, blood pressure, and energy metabolism. The intersection of the two lists (our list and list from [9]) results in 33 genes (Supplementary Table S2), which, according to [9], associated with blood pressure (PDGFB, DDAH1, POMC, REN, SGK1, F2R, NAV2, LRP5, PLCB3), basal metabolic rate (PIK3CD, TG, CCNA1, STK11, PAX8, CDKN1A, CACNA1A), energy metabolism (STK11, DPP4, PLCB3, CACNA1A, GATA4, STX1A, GNB2), response to temperature (ADM, XYLT1, DNAJB6, TRPM2, IL6, MYOF, DNAJC7), smooth muscle contraction (ADM, F2R, PTPRM, KCNMA1, P2RY2), non-shivering thermogenesis (EPAS1, PRDM7, PARK2). Let us further explore more functions of some genes from the intersection.

PIK3CD gene is involved in the development and migration of natural killer cells to inflammation foci and is involved in NK-cell receptor activation [62]. Another gene, IL6, is also found to be associated with the findings of differences in cellular composition between the regions: it is responsible for the differentiation of CD4T cells and takes part in the initiation of the immune response [63]. TG gene is responsible for thyroid hormone thyroglobulin production [64], PAX8 gene is also associated with thyroid function and thyroid-specific gene expression [65]. As shown in [9], the PDGFB gene may be associated with blood pressure regulation. In the context of the difference between the considered regions, the appearance of this gene can be explained by the influence of the ambient temperature on blood pressure, the increase of which is associated with lower temperatures [66]. This may also include the ADM gene, whose activity has also been associated with the production of hypotensive and vasodilator agents [67], as well as the REN gene, which initiates a response cascade to elevate blood pressure [68]. The found gene GATA4, which is included in the regulation of cardiac-specific gene expression, is also associated with cardiac development [69, 70]. Interestingly, the PRDM7 gene itself is presumably associated with epigenetic regulation of gene expression [71].

Indigenous Siberian populations were also studied in [72, 73]. However, there is no overlap between our list of genes and the genes from these works. The reason for this may be that [72, 73] considered a trio of Yakuts-Han Chinese-Europeans, whose differences in adaptation to climate and/or nutritional aspects are much higher than those between Yakutia and Central Russia.

For further GSEA analysis, we used the adaptive methylGSA method [74], which does not use pre-selected lists of CpG sites and genes according to some threshold value, but works with all available CpG sites and their p-values. As a result, methylGSA identified 17 statistically significant terms (with adjusted p-value < 0.05) from the Gene Ontology library [52, 53] (Supplementary Table S4). Among the found terms we can highlight one related to actin filament activity, which regulate cellular behavior and are involved in the aging process and age-associated diseases [75, 76]. Another term related to drinking behavior reflects the mode of water consumption, and the difference between the regions may be related to water supply, water pretreatment, and sanitation problems in Yakutia [77]. The group of terms related to channel activity is especially interesting in the context of differences between Yakutia and Central Russia. The sensation of low temperature through the skin sensory nerve terminals and the propagation of action potentials in cold-sensitive nerve fibers affects a large number of ion channels [78], thus they are closely related to cold susceptibility [79, 80] and may differ in populations living in significantly different climates. The presence of terms concerning the regulation of steroid and corticosteroid hormone secretion is also legitimate, since they are related to endocrine function regulating many aspects of human life, likely playing a role in the developmental processes that lead up to age-specific early life-history transition and thus also in the aging process [81, 82] and adaptation to the cold exposure [83], which are most interesting in the context of our study. Cell fate commitment related to metabolism has also been shown to be an important term in the aging process [84, 85]. Nutritional differences between regions and the increased risk of digestive system diseases in the Yakutian population [86] may be the reason for identifying a term related to digestive system regulation.

It is interesting that among the found terms there are not only those related to adaptation to climatic conditions (water consumption, digestive system regulation), but also those related to aging processes (actin filament activity, cell fate), and both of them (channel activity, regulation of steroid and corticosteroid hormone secretion). Therefore, we further pay attention to aging analysis in the considered cohorts.

#### 2.2.2. Epigenetic age accelerations

To investigate the differences in aging processes between the two regions, we considered 9 types of epigenetic ages: the 4 most common clocks from Horvath’s online calculator [54] (Horvath DNAm age [37], Hannum DNAm age [36], DNAm PhenoAge [39], and GrimAge [38]) and 5 their PC improvements [56]: PCHorvath1, PCHorvath2, PCHannum, PCPhenoAge, PCGrimAge.

Figure 4 shows the correlation between all these epigenetic age estimators with each other and with the chronological age (left column and top row).

**Figure 4.**
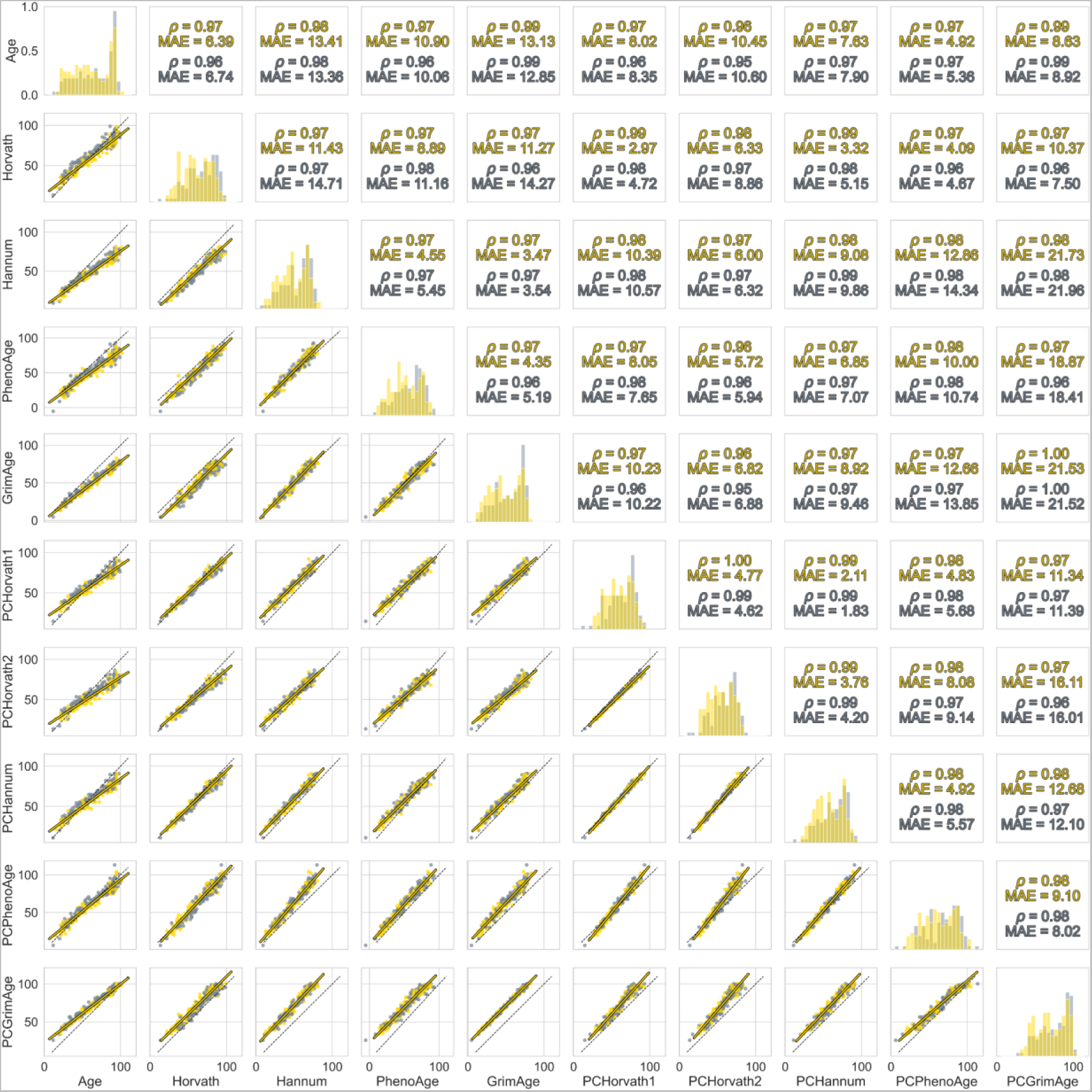
Correlation matrix of chronological age and different types of epigenetic ages. The diagonal elements are overlapping histograms. The bottom triangular elements are scatter plots with the corresponding types of ages for the Central region (yellow) and Yakutia (gray) along the axes. The black dotted line y=x in each scatter plot corresponds to the equality of the plotted ages; the yellow bold line represents the regression plotted on samples from the Central region. The upper triangular elements are the metrics for the corresponding age types: the Pearson correlation coefficient and the mean absolute error for both regions.

The correlation coefficient is close to 1 in all cases, but the MAE is significantly different. The smallest error with chronological age is shown by PCPhenoAge (about 5 years in both regions groups). PCPhenoAge has previously been shown to be more sensitive in predicting the age of hepatocellular carcinoma patients [87] and COVID-19 patients [88]. At the same time, the MAE difference between the regions does not exceed 0.9 years in all cases when comparing chronological age with epigenetic age. The systematic bias and relatively large error between GrimAge and PCGrimAge are also noteworthy, whereas the smallest error (less than 2 years) in both regions is observed between PCHannum and PCHorvath1.

Age acceleration was determined as follows: first, a linear regression was built between the different epigenetic ages and the chronological age for the Central region group (yellow bold lines in Figure 4). Age acceleration values were defined as residuals relative to this linear approximation. Thus, it follows from the definition that in the Central region group, the average age acceleration value is 0, and a non-zero value in the Yakutia group allows us to conclude about accelerated aging in this group relative to the Central region. In addition, the DunedinPACE value also allows us to make conclusions about the differences in age acceleration between regions, since this metric characterizes the pace of aging [57]. Figure 5A-5I shows the distributions of age acceleration values for all considered types of epigenetic clocks with FDR-corrected [51] p-values of Mann-Whitney U Test [89], which reflect the measure of statistical significance of the found differences. Figure 5J shows the DunedinPACE values with the corresponding p-value.

**Figure 5.**
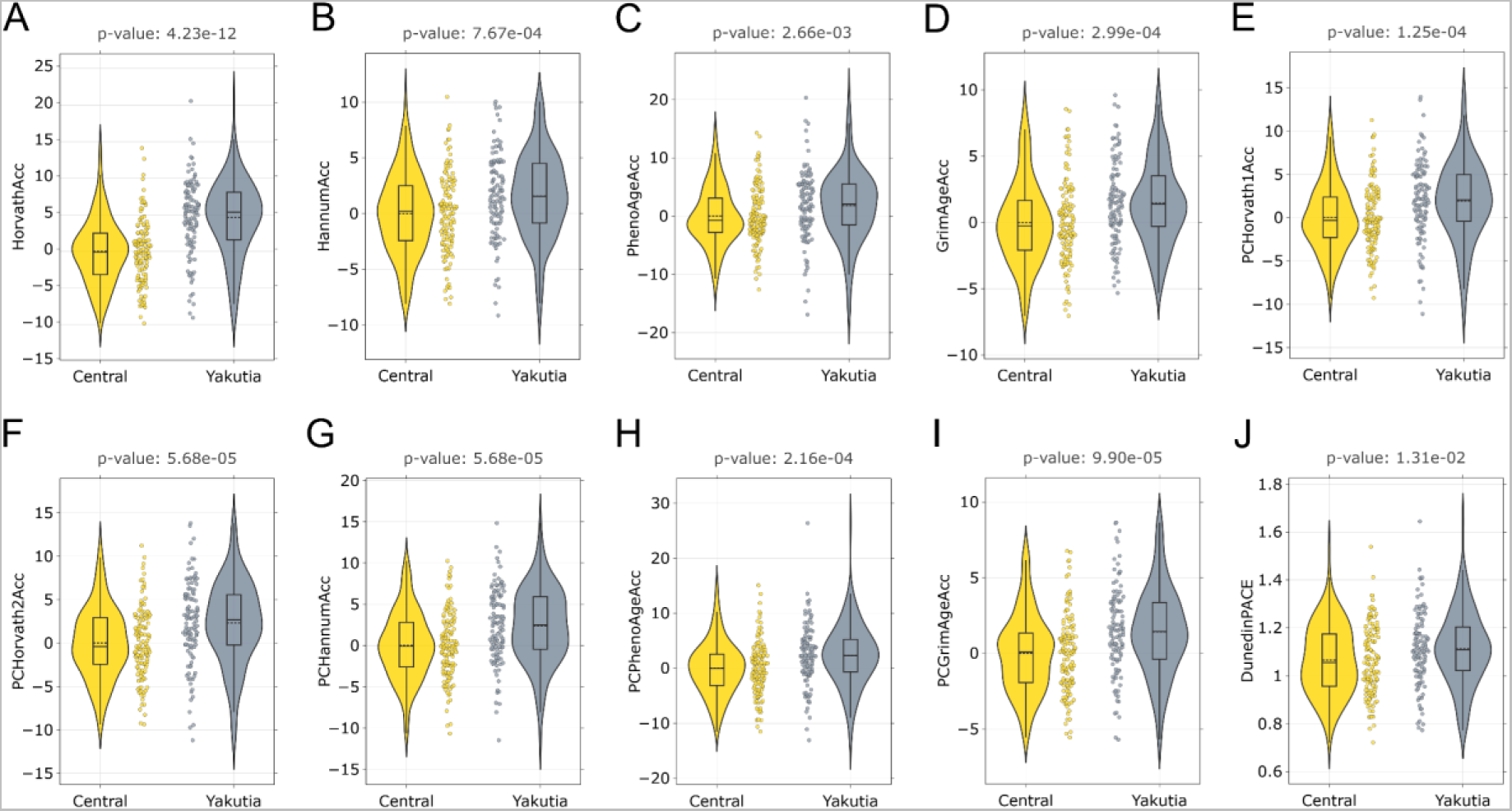
Age acceleration in Central and Yakutia regions for different epigenetic ages: (A) Horvath, (B) Hannum, (C) PhenoAge, (D) GrimAge, (E) PCHorvath1, (F) PCHorvath2, (G) PCHannum, (H) PCPhenoAge, (I) PCGrimAge. Acceleration values were determined as residuals from a linear regression model constructed on samples from the Central region group. Mann-Whitney U test was applied to analyze the statistically significant difference in epigenetic age acceleration between groups. The obtained p-values were adjusted using the Benjamini-Hochberg approach. DunedinPACE values are shown in (J) with the p-values of Mann-Whitney U-test. The solid line in each diagram corresponds to the median value, the dotted line to the mean value.

For all types of epigenetic ages, we observe statistically significant (p-value < 0.05) positive age acceleration in Yakutia relative to the Central region. Horvath DNAm age shows the lowest p-value, and the median acceleration relative to the Central region is 5.36 years. For other epigenetic ages, the median acceleration does not exceed 3 years. It is also interesting that the average DunedinPACE value (Figure 5J) exceeds 1 for both regions, which may indicate that the Russian population ages faster overall than the Dunedin Study participants, whose data were used to develop this metric [57]. However, the same effect for DunedinPACE that was found for all epigenetic clocks persists: in the Yakutia group, the age acceleration is statistically higher with respect to the Central region (median DunedinPACE is 1.11 in Yakutia, 1.05 in Central Russia). Age acceleration in the Yakutia group may also be related to the fact that the original epigenetic clock models were built on data for different populations. To train the Horvath model, 39 datasets were used including participants from different African, Asian, Hispanic, and Caucasian populations. The Hannum model used participants of Caucasian and Hispanic origin. Participants from NHANES for the PhenoAge model were of African American and Caucasian ancestry. GrimAge was built using Caucasian participants from the Framingham heart study Offspring Cohort. For PC clock modifications, data of participants from the London Life Sciences Prospective Population study, of South Asian origin, were used.

#### 2.2.3. Blood cell counts estimation differences

Horvath’s online calculator [54] allows to obtain the distributions of blood cell count measures: CD8T, CD4T, NK, Bcell, Mono, and Gran using Houseman algorithm [55]. The quantitative composition of blood cell populations can vary by race/ethnicity. For example, Tsimane have been shown to have decreased naïve CD4+ T cells and increased exhausted T cells, which may be one of the reasons for their low life expectancy, which was less than 55 years at the beginning of the 20th century [31]. It has also been shown that there is a difference in the levels of naïve CD8+ T cells and naïve CD4+ T cells, as well as different estimated proportions of neutrophils, B-cells and natural killer cells in different races [20, 31].

Figure 6 shows the distribution of blood cell counts for individuals from the Central Russia and Yakutia regions with FDR-corrected [51] p-values of Mann-Whitney U Test [89].

**Figure 6.**
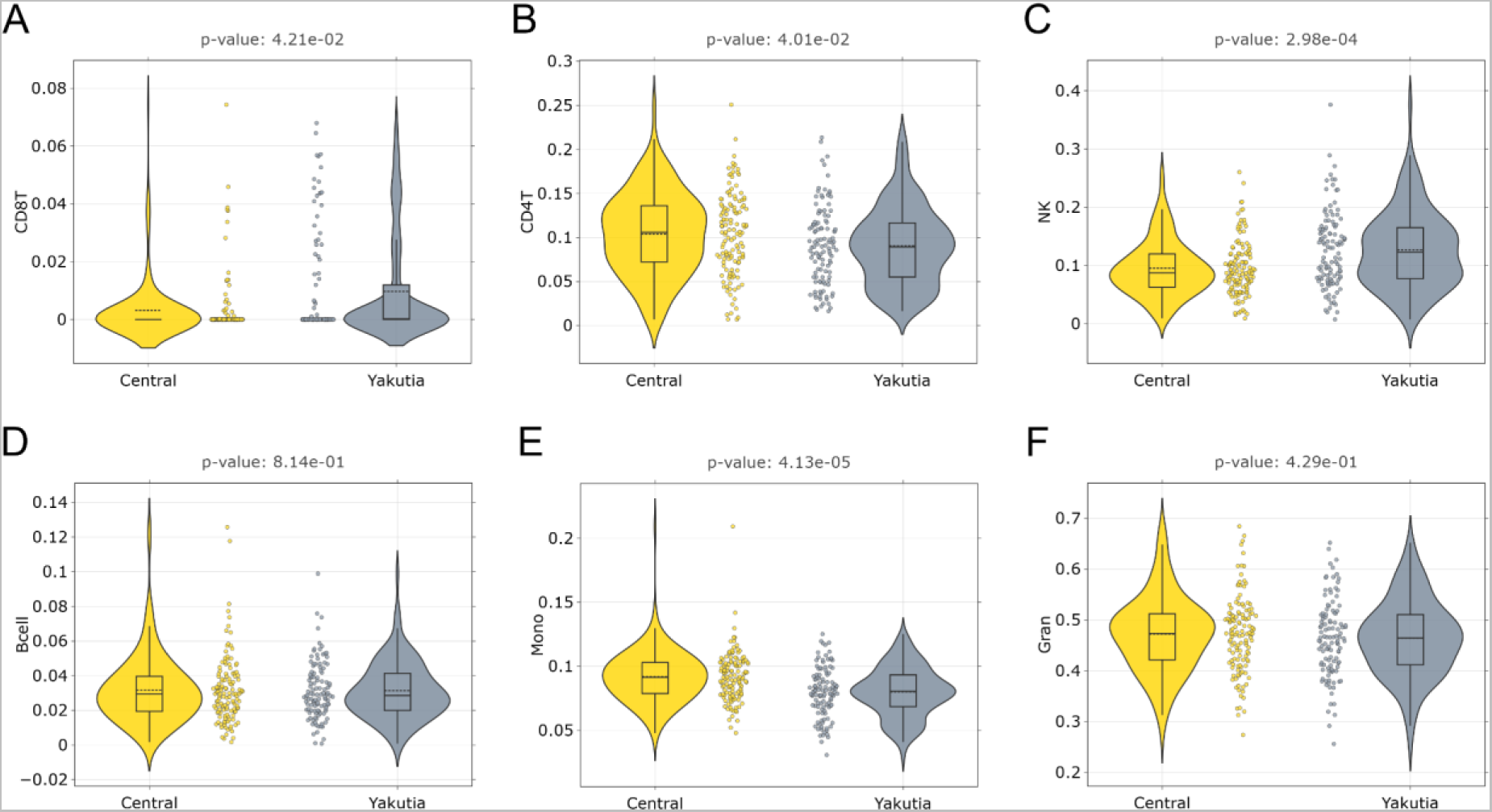
Blood cell count distribution in Central and Yakutia regions: (A) CD8T, (B) CD4T, (C) NK, (D) Bcell, (E) Mono, (F) Gran. Mann-Whitney U test was applied to analyze the statistically significant difference in distribution between the groups. The resulting p-values were FDR-corrected with the Benjamini-Hochberg approach. The solid line on each boxplot corresponds to the median value, dashed line - to mean value.

Statistically significant increased numbers of CD8T and NK are observed in the Yakutia region relative to Central. In contrast, the values of CD4T and Mono are statistically lower in Yakutia than in the Central region. CD8T cells are responsible for the response to the impact of external pathogens unfamiliar to the body [90], their increased number may be associated with an increased viral load in Yakuts compared to the residents of Central Russia. This is consistent with an increase in Natural killer cells, which, along with their main function can produce high levels of cytokines [90]. Differences in CD4T levels, as well as in CD8T and NK levels, may also be due to ethnic differences [91]. For monocytes, European ancestry has been shown to have higher levels of monocytes compared to other populations [92].

### 2.3. Sex-specific differences in regions

This section examines epigenetic differences between males and females independently in the two regions, then compares the results to highlight sex-specific differences between the regions. In the Central region, the sex distribution among participants was 78 females and 53 males, and in Yakutia, 63 females and 51 males (Figure 1B). The sex-specific analyses in each region follow the workflow shown in Figure 1C.

#### 2.3.1. Sex-specific CpGs, DMPs, and GSEA

Further, following the general pipeline (Figure 1C), we studied epigenome-wide differences between males and females in both regions using limma [58]. The resulting distribution of p-values of sex-specific CpGs in both regions (Figure 7A, 7B for Central region and Figure 7C, 7D for Yakutia) differs significantly from the similar distribution in the region-specific task (Figure 2A, 2B): the number of statistically significant CpG sites in both sex-specific tasks is significantly lower than in the region-specific task. However, we should note the similarity in the distributions of sex-specific CpGs between the two regions, which can be observed especially clearly in the p-value and fold change ranges in Figure 7B, 7D. Examples of the distributions of hypo- and hypermethylated sex-specific CpGs for both regions are shown in Figure 8.

**Figure 7.**
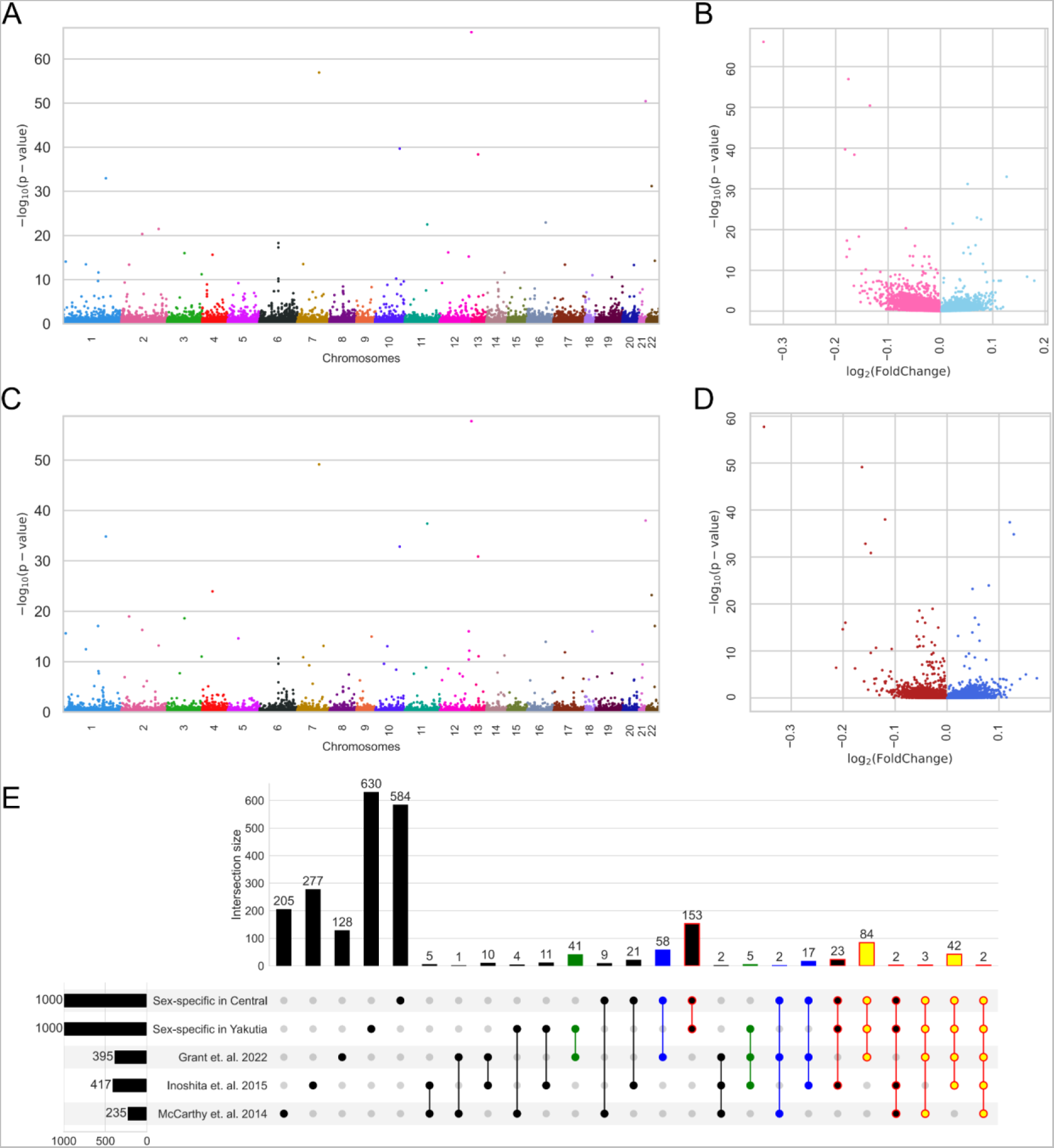
DMP analysis of sex specificity in the Central and Yakutia regions. Manhattan plots for the distribution of adjusted p-values of sex-specific CpGs distributed by location in chromosomes for (A) Central and (B) Yakutia regions; volcano plots of limma results for all CpGs: (C) in the Central region, hypermethylated in females CpGs are highlighted in pink and hypermethylated in males CpGs are highlighted in light blue; (D) in the Yakutia region hypermethylated in females CpGs are highlighted in red and hypermethylated in males CpGs are highlighted in dark blue. Examples of the distributions of differentially methylated CpG by sex in the two regions are shown in Figure 8. (E) UpSet plot showing the intersections of the Top-1000 most statistically significant sex-specific CpGs in Central and Yakutia regions with the lists of sex-specific CpGs from other published studies. Each column corresponds to the intersection of all lists. The sum of all elements in a row is the total value of the elements in the selected CpG list (this number is given on the left barplot: 395 for Grant et. al. [44], 417 for Inoshita, et. al. [93], 235 for McCarthy, et. al.[94], 1000 for both our sex-specific CpG lists). All subsets that include common elements of sex-specific lists for the Central and Yakutia regions from this study are highlighted in red. Yellow indicates subsets of CpGs common to the three sets: sex-specific in Central, sex-specific in Yakutia, and from Grant et. al. [44]. Blue indicates subsets of CpGs common to the two sets: sex-specific in Central and Grant et. al. [44] (without Yakutia). Green indicates subsets of CpGs common to the two sets: sex-specific in Yakutia and Grant et. al. [44] (without Central).

**Figure 8.**
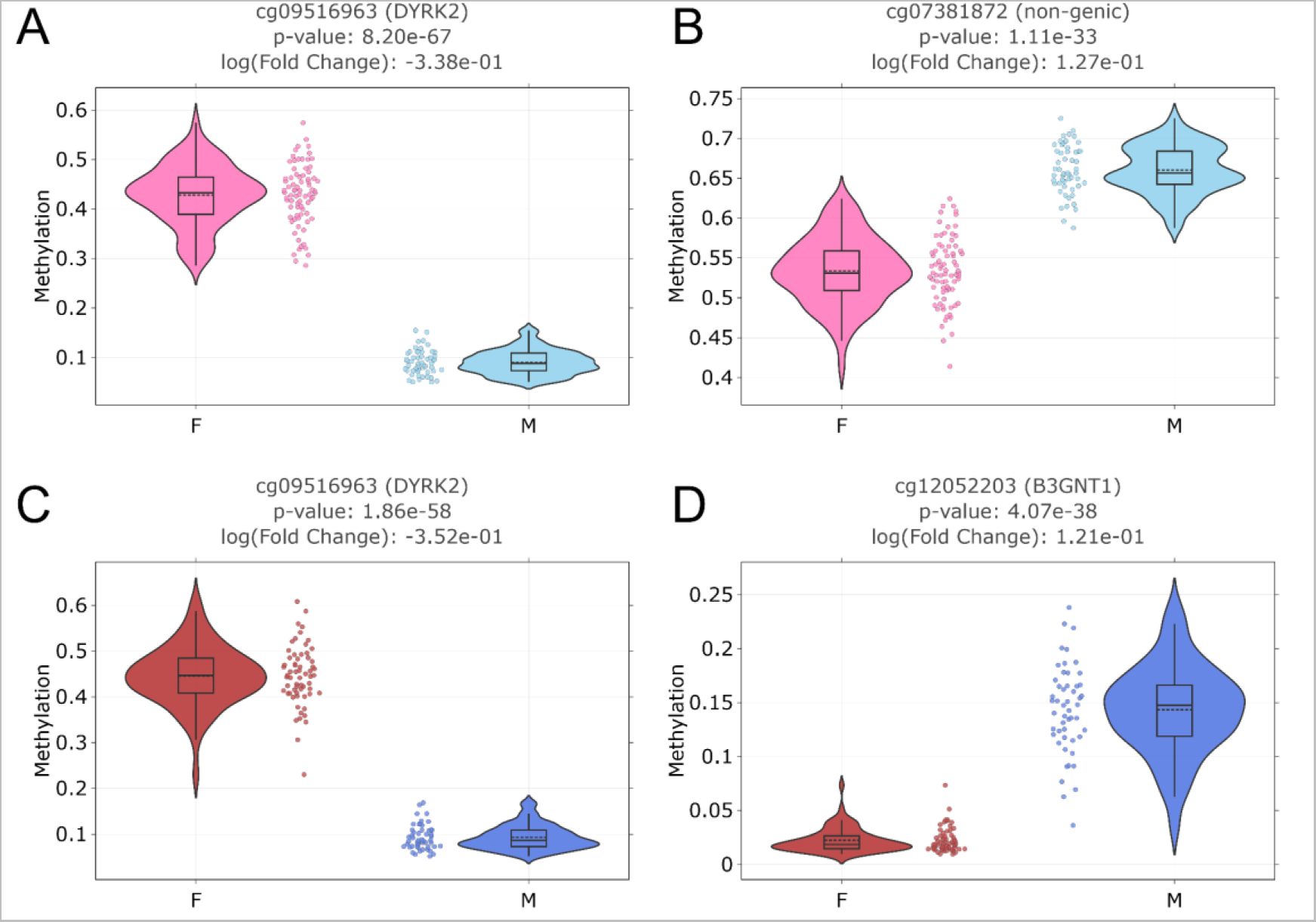
Examples of the sex-specific DMPs distribution in the two regions: (A) hypermethylated in females for Central region; (B) hypermethylated in males for Central region; (C) hypermethylated in females for Yakutia; (D) hypermethylated in males for Yakutia.

Selecting the Top-1000 sex-specific CpGs (Supplementary Table S5) with the lowest p-values in both regions, it appears that these lists overlap by about a third (309 CpGs, which are the sum of all bars with red highlighting in Figure 7E). In addition, we considered the intersections with the lists of sex-specific CpGs from other papers, which also investigated various aspects of sex-specificity in epigenetics. One of the first meta-analyses of autosomal chromosome DNA methylation to identify sex-specific CpG sites was performed in McCarthy, et. al. [94]. 235 CpG sites were detected after correction for multiple testing. Using full-genome DNA methylation profiling, Inoshita, et. al. [93] investigated the effect of sex using multiple linear regression analysis corrected for age and inferred blood cell proportions. 417 CpG sites showed significant gender differences in DNA methylation. One of the most recent studies of sex-specific DNA methylation patterns by Grant, et. al. [44] used data from the Illumina EPIC standard and identified 395 sex-associated CpG sites.

A detailed representation of all possible intersections of the lists of sex-specific CpGs is illustrated in Figure 7E. The largest overlap between the two lists proposed in this paper for Central and Yakutia regions was found with the chronologically most recent work of Grant, et. al. [44]. All possible subsets involving all 3 lists, Central, Yakutia and Grant, et al. are highlighted in yellow (131 CpGs in total). If we consider the lists of the two regions separately, the Central region overlapping with Grant, et. al. gives 208 CpGs (sum of yellow and blue bars on Figure 7E) and for Yakutia region - 177 CpGs (sum of yellow and green bars on Figure 7E).

Thus, the lists of both regions include almost half of all sex-specific CpGs from the work [44]. CpG sites located in noteworthy genes according to [44] were found in the intersection. In particular, CpGs located in the DDX43 gene, which are involved in spermatogenesis and male fertility [94], or CpG located in the GABPA gene, associated with early onset Alzheimer’s disease, Parkinson’s disease, and breast cancer [95]. All the overlaps with other works turned out to be relatively smaller and are also presented in Figure 7E.

The results of the enrichment analysis of the Top-1000 sex-specific CpGs for the both regions are shown in Figure 9. CpGs for the Central region were overrepresented in the 6th chromosome, and for Yakutia in the 4th chromosome. Interestingly, the work [44] also observed the largest number of sex-specific CpGs in chromosome 6 (this work considers the population from the UK, which is just more similar in terms of ethnicity to Central Russia than to Yakutia). Statistically significant underrepresentation in the gene body is observed in both lists. Also, for both lists there is overrepresentation in high-density CpG islands and shores and underrepresentation in low-density OpenSea regions.

**Figure 9.**
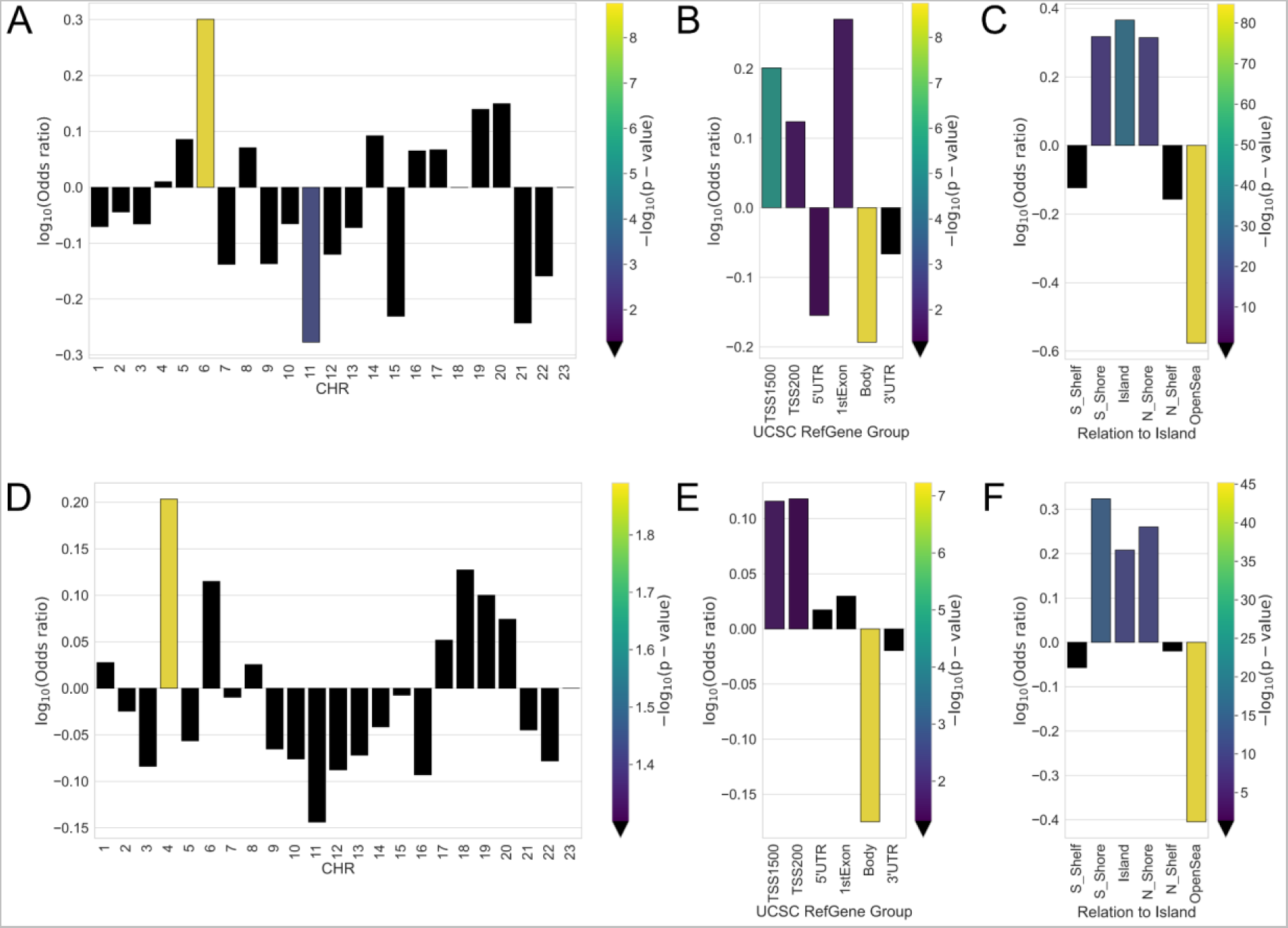
Enrichment for the Top-1000 sex-specific CpGs in Central region of: (A) chromosomes, (B) genomic region, (C) relation to CpG island; In Yakutia of: (D) chromosomes, (E) genomic region, (F) relation to CpG island. Odds ratio values and corresponding p-values (shown by color) were obtained from Fisher exact test. Black color indicates the absence of statistical significance (p-value > 0.05).

Further GSEA analysis, performed independently for both regions, revealed 40 terms in the GO library for the Central region and 39 terms for Yakutia (Supplementary Table S6). Moreover, the lists of terms for the two regions are almost identical: 39 common terms and only 1 specific for the Central region (GO:0004713 protein tyrosine kinase activity). Among the common terms appear ones related to fat cell proliferation, whose differences between the sexes have been well studied [96, 97]. Next, a whole group of terms related to the processes of glycogen, glucose, and carbohydrate metabolism is worth mentioning. The processes associated with glycogen metabolism during exercise differ significantly in men and women, with less muscle glycogen being depleted in women [98, 99]. In the resulting list, 6 of 40 terms are related to glycogen. The processes of glucose metabolism are also related to it. In women, glucose metabolism is affected by the menstrual phase, as well as glucose production in the liver is lower and glucose appearance rates are higher [99]. Because of this, the incidence of different types of diabetes differs in men and women [100]. Terms related to carbohydrate processes contribute to 10% of the total number of sex-specific terms. Women tend to oxidize less total carbohydrates than men in response to physical activity [98, 101]. The term manganese ion binding may reflect the fact that men in general absorb less manganese than women [102].

#### 2.3.2. Epigenetic metrics: age accelerations and blood cell counts

Similar to the analysis of between-region differences (Section 2.2), 9 types of epigenetic ages (4 types of classical clocks [36–39] and their PC variations [56]), DunedinPACE values [57], and blood cell counts measures [55] were considered.

We investigated the correlation of all epigenetic ages with each other along with chronological age separately for males and females in the two regions. The two regions display differences in MAE values between males and females when comparing chronological ages with epigenetic ages. Hannum, Grim, and PCGrim ages tend to have the same error difference between the sexes in both regions: Hannum age has an error for females about 2-3 years larger than for males, GrimAge has an error for females about 4-5 years larger than for males, and PCGrimAge, by contrast, gives about 3-4 years less error for females in both regions. However, there are differences between regions: for example, Horvath age has a larger error for females in the Central region and for males in the Yakutia region (relative to chronological age). On the whole, if we consider the correlations of different ages and their MAE relative to each other in males and females in both regions, similar trends are observed.

Age acceleration within each region was determined as follows: a linear regression was built between chronological age and corresponding epigenetic age on the female group only, and the acceleration values were derived from these linear models. As a result, in both regions, the mean acceleration value for the female group will be 0 (by construction), while for the male group a non-zero value will be obtained, based on which we can conclude about sex-specific age acceleration in the two regions. The difference in the distribution of age acceleration values between males and females was tested using the Mann-Whitney U Test with FDR-corrected p-values < 0.05. An additional characteristic that allows us to make conclusions about sex-specific age acceleration in the two regions is DunedinPACE with the corresponding p-value. Figure 10 shows the summary of statistically significantly different epigenetic biomarkers between males and females in both regions.

**Figure 10.**
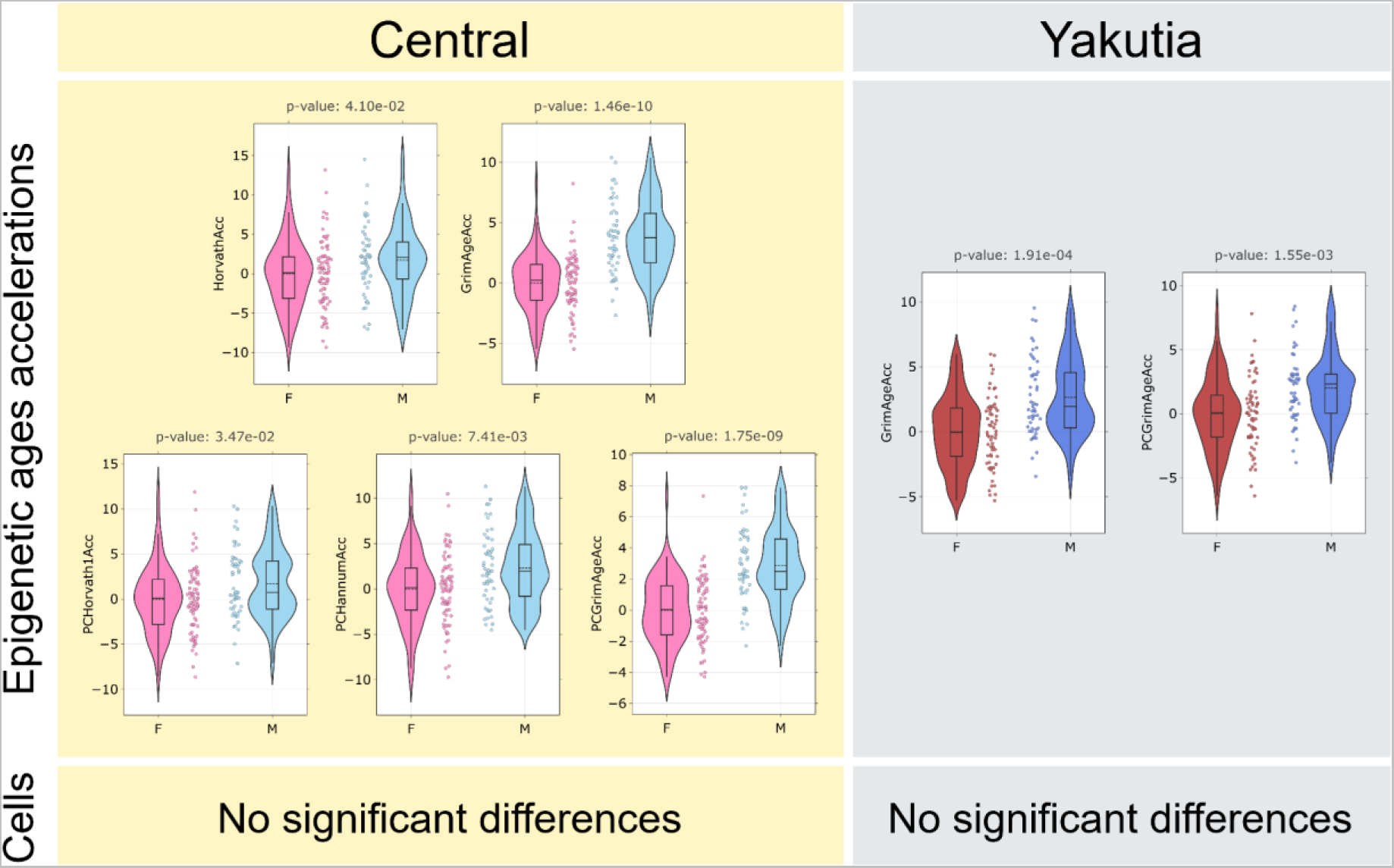
Summary of statistically significantly different epigenetic biomarkers between males and females in the Central region (left yellow column) and Yakutia (right gray column). Mann-Whitney U-test was applied to analyze the statistically significant difference in the distribution between the groups separately for all epigenetic ages and for cell counts. The obtained p-values were corrected using the Benjamini-Hochberg method. Only significant biomarkers are shown. The solid line in each diagram corresponds to the median value, the dotted line to the mean value.

In the Central region, a positive statistically significant age acceleration in the male group was found in 5 of the 9 types of epigenetic ages, while in Yakutia it was found only for 2 types, and they are GrimAge and its PC-modification, which are also statistically significant in the Central region. Several studies have shown that DNAm GrimAge shows more accurate results compared to other models, as well as confirming the thesis of slower aging rates in females compared to males [103–105]. The distribution of DunedinPACE values is not statistically different between males and females in different regions, nor are blood cell counts estimations.

## 3. Discussion

### 3.1. Conclusion

This work is the first epigenetic study focused on the Yakutian population that try to elucidate different dimensions of the variability observed addressing: 1) epigenetic features of the adaptation mechanisms to the cold; 2) pattern of age acceleration and aging rate in a population characterized by extreme environmental conditions and 3) sex-specific methylation patterns. In this work, we examined the epigenetic features of the 114 individuals from the Yakutian population in comparison to 131 participants from Central Russia. The Yakutian population live in the extreme environment and experience an extended period of severe cold temperatures, dramatic variation in photoperiod and variability in food resources. Participants from Central Russia live in a milder environment, with higher temperatures in a shorter winter period compared to Yakutia. The considered populations also differ in terms of genetic history: representatives of Central Russia are close to the peoples of Northeast Europe, while Yakutia shows genetic components that characterized Siberian populations and East Asian populations. First, we focused specifically on region-specific differences between the considered regions, and next, we studied sex-specific differences in both regions and compared them.

We selected 1000 most region-specific CpGs, for which enrichment analysis showed statistically significant overrepresentation in chromosome 16, in the regulatory genomic region TSS1500 and in CpG islands. We intersected the list of genes we obtained from the list of the most statistically significantly different CpGs with the list from Cardona, et. al. [9], a notable work analyzing single-nucleotide polymorphism data from Siberian populations, including Yakutian, and identifying genes associated with biological processes hypothetically associated with adaptation to the cold. 33 genes appeared in the intersection, most of which are associated with blood pressure, basal metabolic rate, energy metabolism and response to temperature, suggesting that part of the epigenetic variability observed could be ascribable to genetic background that characterized this population.

Among the overlapping genes, there are also those whose functions are closely related to immune responses and to the considered blood cells, such as NK cells and CD4T cells (PIK3CD and IL6 genes). Interesting correlation is found between the TG gene responsible for thyroid hormone thyroglobulin production [64] and the terms related to endocrine system functioning and hormone production found by GSEA. Glucocorticoids regulate thyroid function, thereby influencing the production of thyroglobulin [106]. The KCNMA1 gene, which is associated with the regulation of the contraction of smooth muscle, also has channel activity among its functions, which appeared among important GO terms [107, 108]. CACNA1A [109] and STX1A [110] genes are also associated with channel activity.

GSEA of statistically significantly different CpGs by methylation levels between the two regions revealed 17 Gene Ontology terms. Among the found terms we can highlight the ones related to actin filament activity, channel activity, regulation of steroid and corticosteroid hormone secretion, as well as a term related to regulation of the digestive system. Among the found terms, there are those related to adaptation processes to the cold climate, to aging processes, and to both aspects.

To analyze the aging process, we considered the 4 most common types of epigenetic ages (Horvath DNAm age, Hannum DNAm age, DNAm PhenoAge, and GrimAge) and their 5 PC modifications (PCHorvath1, PCHorvath2, PCHannum, PCPhenoAge, PCGrimAge). All these metrics were highly correlated with each other, with PCPhenoAge showing the lowest MAE. When comparing the age acceleration of the studied models and DunedinPACE aging rate, we found that in Yakutia, for all considered epigenetic ages, there is a statistically significant positive age acceleration compared to Central region representatives. At the same time, the average DunedinPACE aging rate exceeds 1 in both regions (in other words, the Russian population as a whole is aging accelerated, with more than one biological year per chronological year). All considered epigenetic models are based on data from participants of different populations mainly of European ancestry (mostly industrialized ones), without including the indigenous populations, which may affect the results. Epigenetic age acceleration in Yakutia and increased aging rates in both considered populations may signal the need for increased attention to healthcare and public health problems.

Differences between regions are also observed for blood cell composition: statistically significant increased number of CD8T and NK and statistically significant decreased number of CD4T and Mono are observed for the Yakutia region.

Next, we analyzed sex-specific differences in the two regions and selected 1000 most sex-specific CpGs in each region. We also considered the lists of sex-specific CpGs from Grant, et. al. [44], McCarthy, et. al. [94], Inoshita, et. al. [93] and compared them with the lists obtained for the Central and Yakutia regions. The largest overlap of our lists is obtained with [44] (almost half of the list), and CpGs corresponding to genes related to spermatogenesis and various diseases can be found in the overlap.

GSEA of statistically significantly different CpGs by methylation levels between the sexes in both regions revealed almost identical Gene Ontology terms for the Central region and Yakutia: 39 common terms and one specific term for the Central region. Among the found terms we can highlight the ones related to the processes of glycogen, glucan, glucose, and carbohydrate metabolism, as well as fat cell proliferation and manganese ion binding. However, we did not find clear region-specific differences between the sexes in the highlighted biological functions. As for regional differences, we next focused on the analysis of sex-specific epigenetic differences in the aging process in the two regions. We considered 9 types of epigenetic clock (classical ones and PC modifications), DunedinPACE metric, and blood cell counts. Correlations and MAE for the epigenetic ages considered have similar trends for both regions in men and women: correlation coefficients in males and females are similar, but most models show a higher error in females than in males when compared to chronological age. 5 out of 9 epigenetic age models showed statistically significant positive age acceleration in men compared to women in the Central region, while in the Yakutia region only GrimAge with its PC modification showed the same result. At the same time, neither DunedinPACE, nor blood cell counts had statistically significant differences between males and females in both regions. Thus, we performed the first study of the epigenetic data of the Yakutian cohort, paying special attention to region-specific features, aging processes, age acceleration, and sex-specificity. We revealed geographic region-specific differentially methylated regions associated with both, adaptation to climatic conditions and aging processes. We also showed that representatives of the Yakutia region show higher age acceleration compared to Central Russia, one of the reasons for which may be the more severe climatic conditions in which Yakuts live and the need to adapt to the cold. Other possible factors that we do not take into account in this study may include diet, pathogen load, socioeconomic conditions, and many others. However, a certain degree of age acceleration is found for Central Russia too. For both regions, we confirmed that men age faster than women (resulting in a shorter life expectancy), but no significant sex-specific difference was found between the regions.

### 3.2. Limitations

We would also like to address the limitations of this work. The most common limitation of many studies investigating biomedical data is sample size. Data such as DNA methylation often have small sample sizes compared to their dimensionality. Another limitation is the wide variety of methods available for performing GWAS and EWAS analyses, which take into account different background information, require a selection of thresholds, and can lead to varying results. In terms of interpretation of the obtained results, differences between regions can be caused by both genetics and environment, and it is difficult to disentangle their influence.

## 4. Methods

### 4.1. Data collection

All study participants were explained the specifics of the procedure, possible inconveniences and risks. Each participant signed an informed consent and filled out a consent for personal data processing, taking into account the principle of confidentiality (accessibility only to the research group and presentation of data in a common array). The study was approved by the local ethical committee of Nizhny Novgorod State University. All research procedures were in accordance with the 1964 Helsinki Declaration and its later amendments.

All study participants were healthy; exclusion criteria included chronic diseases in the acute stage, cancer and acute respiratory viral infections at the moment of biomaterial donation, as well as pregnancy in women.

The main criterion for inclusion of participants from Yakutia in this study was ethnicity. All participants are Yakuts (Sakha) in three generations. They were born and live on the territory of the Republic of Yakutia (Sakha) and their parents and ancestors are indigenous to Yakutia. All participants from the Central region are native Russian residents.

### 4.2. DNAm processing

Phenol Chloroform DNA extraction was used. DNA was quantified using the DNA Quantitation Kit Qubit dsDNA BR Assay (Thermo Fisher Scientific) and 250 ng was bisulfite-treated using the EpiMark Bisulfite Conversion Kit (NEB) with case and control samples randomly distributed across arrays. The Illumina Infinium MethylationEPIC BeadChip [111] was used according to the manufacturer’s instructions. DNA methylation is expressed as β values, ranging from 0 for unmethylated to 1 representing complete methylation for each probe. DNAm data preprocessing, normalization, and batch effect correction were performed with the standard pipeline in the ChAMP [112, 113] R package. The preprocessing was as follows: (1) Probes with a detection p-value above 0.01 in at least 10% of samples were removed; (2) Probes with a beadcount less than three in at least 5% of samples were removed; (3) All non-CpG probes [114], SNP-related probes [115], and multi-hit probes were removed [116]; (4) All probes located on chromosomes X and Y were filtered out. It is also worth noting that all remaining samples have less than 10% of probes with a detection p-value above 0.01 and they don’t need to be excluded. As a result, 739,168 СpGs remained for the analysis. Functional normalization of raw methylation data was performed using minfi [117] R package function. The ComBat method [118, 119] was used to correct for Slide and Array batch effects.

### 4.3. DMPs

DMPs analysis was performed using limma method [58] generalization in the ChAMP [112, 113] R package. Region and sex were used as categorical variables to perform a contrast comparison between the two phenotypes. This method provides adjusted p-values [51] of the statistical test for the difference in methylation levels between the two considered groups, as well as fold change, which indicates the differences in mean values between the groups.

### 4.4. GSEA

Traditional approaches to testing a gene set can produce biased results because of differences in gene length, and the number of CpG sites can vary even among genes of the same length. EWAS can lead to multiple p-value associations per gene, so it is necessary to consider the number of CpGs instead of gene length, and considering all p-values provides an opportunity to take into account dependencies between genes in the set. MethylGSA R package [74] takes all these details into account, therefore we used this method. It allows testing of gene sets with length bias adjustment in DNA methylation data. Enrichment of GO annotations was calculated using the methylglm function, which takes p-values of each CpG and implements logistic regression adjusted for the number of probes in the enrichment analysis. The minimum and maximum number of genes in gene sets were set to 10 and 1000, respectively. All other settings were defaults. The result contains gene sets ranked by p-values. GO terms whose adjusted p-values were less than 0.05 were considered statistically significant.

### 4.5. Epigenetic ages and estimates of blood cell counts

We used Horvath’s online calculator [54] to obtain 4 epigenetic ages (Horvath DNAm age [37], Hannum DNAm age [36], DNAm PhenoAge [39], and GrimAge [38]). The Hannum DNAm Age model measures the rate of human methylome aging under the influence of sex and genetic variants with application to different human tissues and is able to highlight certain components of the aging process. The Horvath DNAm Age model can be applied to a wide range of tissues and cell types, allowing to compare the age of different tissues of the same person to identify signs of accelerated aging associated with different diseases. Using a two-step process involving incorporation of clinical measures of phenotypic age, the epigenetic aging biomarker DNAm PhenoAge was developed, and it correlates with age in all tissues and cells tested. This model is capable of capturing the risks of a variety of outcomes in different tissues and cells and highlighting the aging pathways. Another predictor of longevity, DNAm GrimAge, is a biomarker based on a limited number of DNAm surrogates that can predict time to death, heart attack, cancer and other age-associated diseases.

The PC modifications of the epigenetic ages were calculated according to the original algorithm [56]. These PC clocks can minimize noisy PCs, separate noise from age-associated signals and they use information from multiple CpGs, diluting noise from individual CpGs.

To compare different epigenetic ages we used Pearson correlation coefficient [120] and mead absolute error (MAE), calculator by the formula: 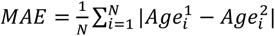, where 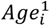 – first compared age, 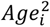 – second compared age, *N* – number of samples.

Age acceleration was determined as follows for regions and sexes. To compare the regions, first, a linear regression was built between the different epigenetic ages and the chronological age for the Central region group. Age acceleration values were defined as residuals relative to this linear approximation. To compare the sexes, a linear regression was built for the females group.

Horvath’s online calculator also allows to obtain the distributions of blood cell counts: CD8T, CD4T, NK, Bcell, Mono, and Gran using Houseman algorithm [55].

The distributions of age-accelerated values and blood cell composition between the groups under consideration were tested using the Mann-Whitney U Test [89]. This is a non-parametric test for comparing results between two independent groups, which is used to test the probability that two samples come from the same population, with a two-sided null hypothesis that the two groups are not the same. All resultant p-values were adjusted according to the Benjamini-Hochberg procedure [51].

Additionally, it is worth mentioning that both the original versions of the epigenetic clock in the Horvath’s calculator and their PC modifications were developed for the Illumina 450k methylation data standard, not for Illumina EPIC, so the values of some CpGs may be missing. Also, some CpGs involved in the calculation of epigenetic clock values are excluded at the data preprocessing stages (filtering, quality control, exclusion of SNP-related probes); therefore, all values of such missing for various reasons CpGs were automatically imputed according to the algorithms in the corresponding articles.

## Data availability statement

DNA methylation profiles generated in this study will be available as the GEO public repository prior to publication.

## Code availability statement

No new or unpublished methods were used in the study. The scripts used in the generation of the manuscript are available on GitHub: https://github.com/GillianGrayson/EWAS_Yakutia.

## Author’s Contributions

Conceptualization: A.Ka., I.Y., E.Ko., M.G.B., C.G., M.V., C.F., M.I.; Investigation: A.Ka., I.Y., E.Ko., R.Z., T.S., M.V., C.F., M.I.; Resources: E.Ko., R.Z., T.S., S.S., A.Ks., M.N., M.V.; Formal analysis: A.Ka., I.Y.; Methodology: A.Ka., I.Y., M.G.B., C.G., M.I.; Software: A.Ka., I.Y.; Supervision: M.G.B., C.G., E. Kh., M.V., C.F., M.I.; Visualization: A.Ka., I.Y.; Writing – original draft: A.Ka., I.Y.; Writing – review & editing: A.Ka., I.Y., E.Ko., M.G.B., C.G., E.Kh., C.F., M.I.

## Abbreviations

DMP: Differentially Methylated Position
DNAm: DNA methylation
EWAS: Epigenome-Wide Association Study
FDR: False Discovery Rate
GO: Gene Ontology
GSEA: Gene Set Enrichment Analysis
GWAS: Genome-Wide Association Study
MAE: Mean Absolute Error
PC: Principal Component
RBC: Red Blood Cell

## Acknowledgements

The authors acknowledge Tatyana Klimova and Vladimir Osakovsky for help in organizing and participation in the discussion.

